# HiCHub: A Network-Based Approach to Identify Domains of Differential Interactions from 3D Genome Data

**DOI:** 10.1101/2022.04.16.488566

**Authors:** Xiang Li, Shuang Yuan, Shaoqi Zhu, Hai-Hui Xue, Weiqun Peng

## Abstract

Chromatin architecture is important for gene regulation. Existing algorithms for the identification of interactions changes focus on loops between focal loci. Here we develop a network-based algorithm HiCHub to detect chromatin interaction changes at larger scales. It identifies clusters of genomic elements in physical proximity in one state that exhibit concurrent decreases in interaction among them in the opposite state. The hubs exhibit concordant changes in chromatin state and expression changes, supporting their biological significance. HiCHub works well with data of limited sequencing coverage and facilitates the integration of the one-dimensional epigenetic landscape onto the chromatin architecture. HiCHub provides an approach for finding extended architectural changes and contributes to the connection with transcriptional output. HiCHub is freely available at https://github.com/WeiqunPengLab/HiCHub.

## Introduction

The three-dimensional (3D) conformation of chromatin structure is closely involved in gene expression and cellular homeostasis ^1^. Spatial contacts between cis-regulatory elements such as enhancers and promoters constitute an important mechanism in transcriptional regulation. Dynamic changes of the 3D organization are associated with cellular processes and experimental perturbation^2, 3, 4^. Alteration of chromatin organization has been implicated in dysregulation, developmental disorders and tumorigenesis^5, 6^. Given the importance of the dynamic changes in chromatin architecture, algorithms have been developed to identify these changes. However, existing algorithms^7, 8, 9, 10, 11^ focus on the changes of punctate interaction between two focal loci. Methodologies for the detection of concurrent changes on larger length scales and/or multiple loci have been underexplored. Recent advances support the biological relevance of this type of changes. Enhancers can cluster along the genome to form super-enhancers (SEs)^12^. which recruit a high level of transcriptional regulators and drive the expression of genes important for cell-type specification. The SE components assemble in close spatial proximity^13^. More recently, spatial clustering of enhancers has been observed in various systems^14, 15, 16, 17^. These enhancer communities were capable of mutually enhancing transcription factor binding and controlling lineage-determining genes^15^. Motivated by these experimental observations, we developed a network-based algorithm that systematically identifies spatial clusters of genomic elements that exhibit concordant directional interaction changes among them from Hi-C data or its variants. We validated identified differential hubs using orthogonal functional genomic and epigenomic data, and found the hubs exhibited concordant changes in chromatin state and transcriptional output.

## Result

### Overview of HiCHub

HiCHub is based on the following hypothesis: a spatial cluster of genomic elements whose interaction among them are subject to changes in the same direction (i.e., increase or decrease) between two cell states is likely to play a significant role in cell-state-specific gene regulation. HiCHub identifies the above configuration by applying a network-based approach to the genome architecture derived from data generated by high-throughput sequencing protocols based on chromatin conformation capture^18^, including Hi-C or its variants such as Micro-C^19^. The schematic workflow of HiCHub is shown in Fig.1. Briefly, in pre-processing the contact matrices of two cell states, *M*_*A*_ and *M*_*B*_, were subject to MA normalization to remove library-specific biases^7^ (Fig. S1). The state A-specific hubs were identified by collecting elements in *M*_*A*_ that decreased in *M*_*B*_ for network construction. The nodes represented the interaction anchors, the edges represented the interactions, and the edge weights were set by the interaction strength in state A. We then used the community_multilevel algorithm in the igraph platform^20^ to find network clusters. These communities of genomics elements tended to be in physical proximity in state A with mutual interaction strengths decreasing consistently in state B. The nodes of a network community are then projected back onto the genome and stitched according to proximity along the genome into candidate hub-anchors. The candidate hubs were defined as the sub-matrices of interactions of one or two candidate hub-anchors, shown respectively as the highlighted triangles and rectangles in step 5 of Fig. 1. The significance of a candidate hub was assessed by evaluating the consistency of the change in interactions in the hub in the desired direction using the Wilcoxon signed-rank test. The state B-specific hubs were identified in a similar manner. The details of the HiCHub algorithm can be found in the Method section.

**Figure 1.**
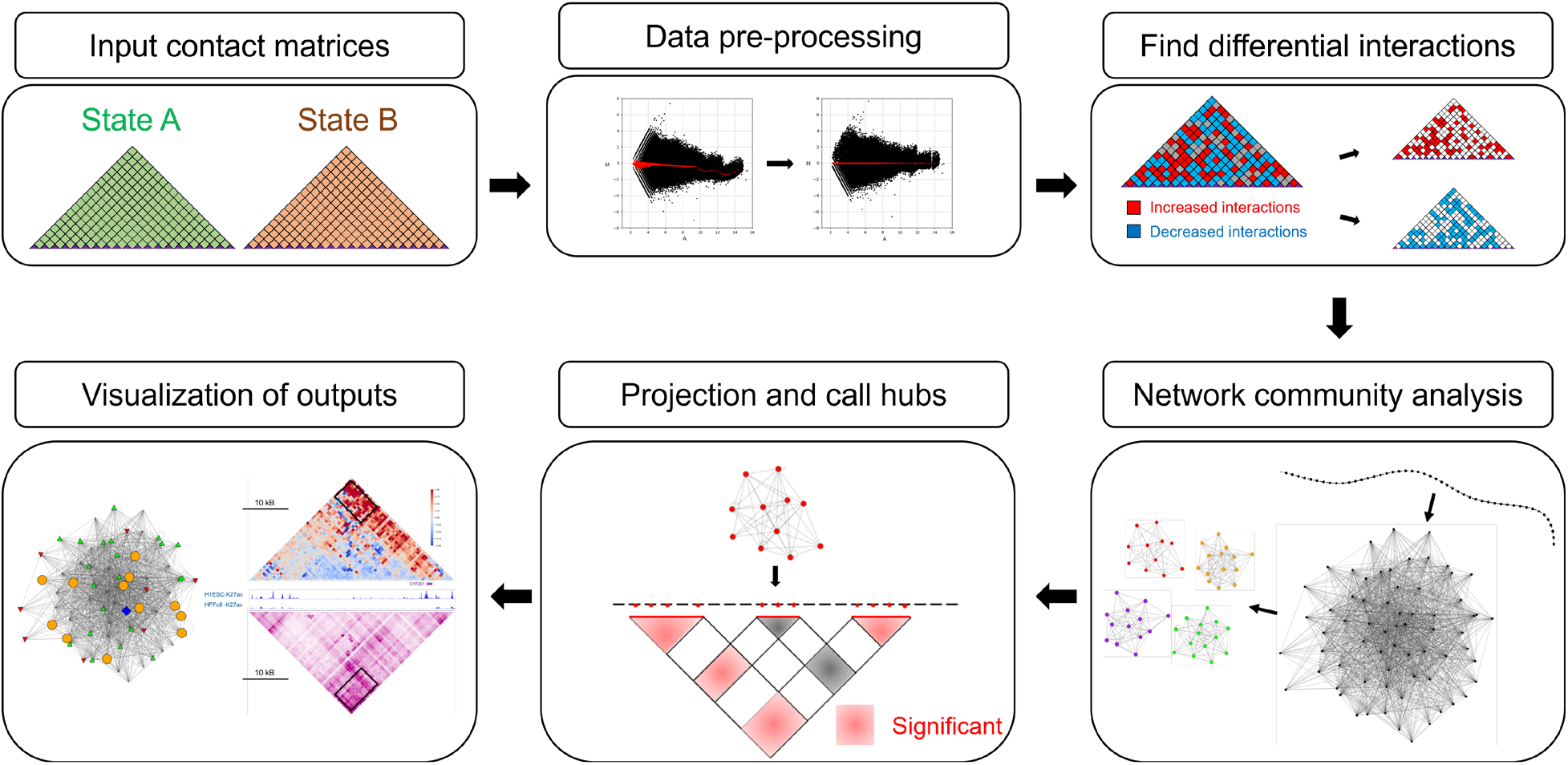
Schematic workflow of HiCHub. 1) Contact matrices with the same resolution serve as input. 2) Data preprocessing involves filtering matrix elements and using MA normalization to balance the matrices. 3) Interactions changing in the desired directions were collected and were used for network construction. 4) Network communities were identified using the community_multilevel algorithm in the igraph platform. 5) The nodes of each community were projected back onto the genome and anchors in proximity along the genome were stitched together forming the anchors of candidate hubs. 6) The significance of candidate hubs (solid triangles and rectangles) was assessed by the consistency of the hub interactions changes between the two states. The red and grey colors denote hubs above and below the significance threshold, respectively. 7)Visualization of HiCHub output. The left panel represents the network community of a hub annotated with epigenomic information. The right panel shows the matrix of interaction differences (top) and the interaction matrix of the reference state (bottom). The hub is highlighted with a blue box.

### Chromatin state and gene expression associated with hubs exhibit changes concordant with interaction changes

To evaluate its performance, we applied HiCHub to a set of Micro-C data, which mapped the chromosome architecture in human embryonic stem cells (H1-ESC) and human foreskin fibroblast cells (HFFc6)^19^ (Table S1). Using contact matrices with a resolution of 10 kilobases (Kb) and a stringent significance threshold of *p*-value < 1E-20, we detected 1930 hubs with higher interaction in H1-ESC cells (i.e., H1-ESC-specific) and 734 hubs with higher interaction in HFFc6 cells (i.e., HFFc6-specific). The median size of the hub-anchors was about 180 kilobases (Kb), significantly larger than that of a typical loop anchor. The reproducibility of hubs was evaluated using individual bio-replicates (Fig. S2).

CTCF is known as a master weaver of the 3D genome^21, 22^. It is involved in both architectural and regulatory chromatin interactions^23, 24 25^ and mediates extensive cell-type-specific interactions^26^. Therefore, we reasoned that the extensive interaction changes on hubs could be associated with corresponding changes in CTCF binding. By comparing CTCF binding intensity on CTCF binding sites in both cell-types downloaded from the ENCODE portal^27, 28^ (Table S1), we found CTCF exhibited stronger binding in H1-ESC cells than in HFFc6 cells on anchors of H1-ESC-specific hubs, and vice versa on HFFc6-specific hubs (Fig. 2a). We then evaluated the differences in regulatory activities on the differential hubs. Open chromatin regions harbor regulatory elements. We found that chromatin accessibility on open chromatin regions measured by DNase-Seq^27, 28^ (Table S1) exhibited concordant changes on hub anchors. Chromatin accessibility was higher in the H1-ESC cells than in the HFFc6 cells on anchors of H1-ESC-specific hubs, whereas HFFc6-specific hubs exhibited the opposite trend (Fig. 2b). Histone modification H3K27ac is associated with the activity of the enhancers and promoters. H3K27ac levels on its peak regions^27, 28^ (Table S1) on hubs exhibited similar behavior (Fig. 2c) across the two states. Finally, we asked whether the significant differences in interaction changes and chromatin state changes on the differential hubs lead to differences in gene expressions.

**Figure 2.**
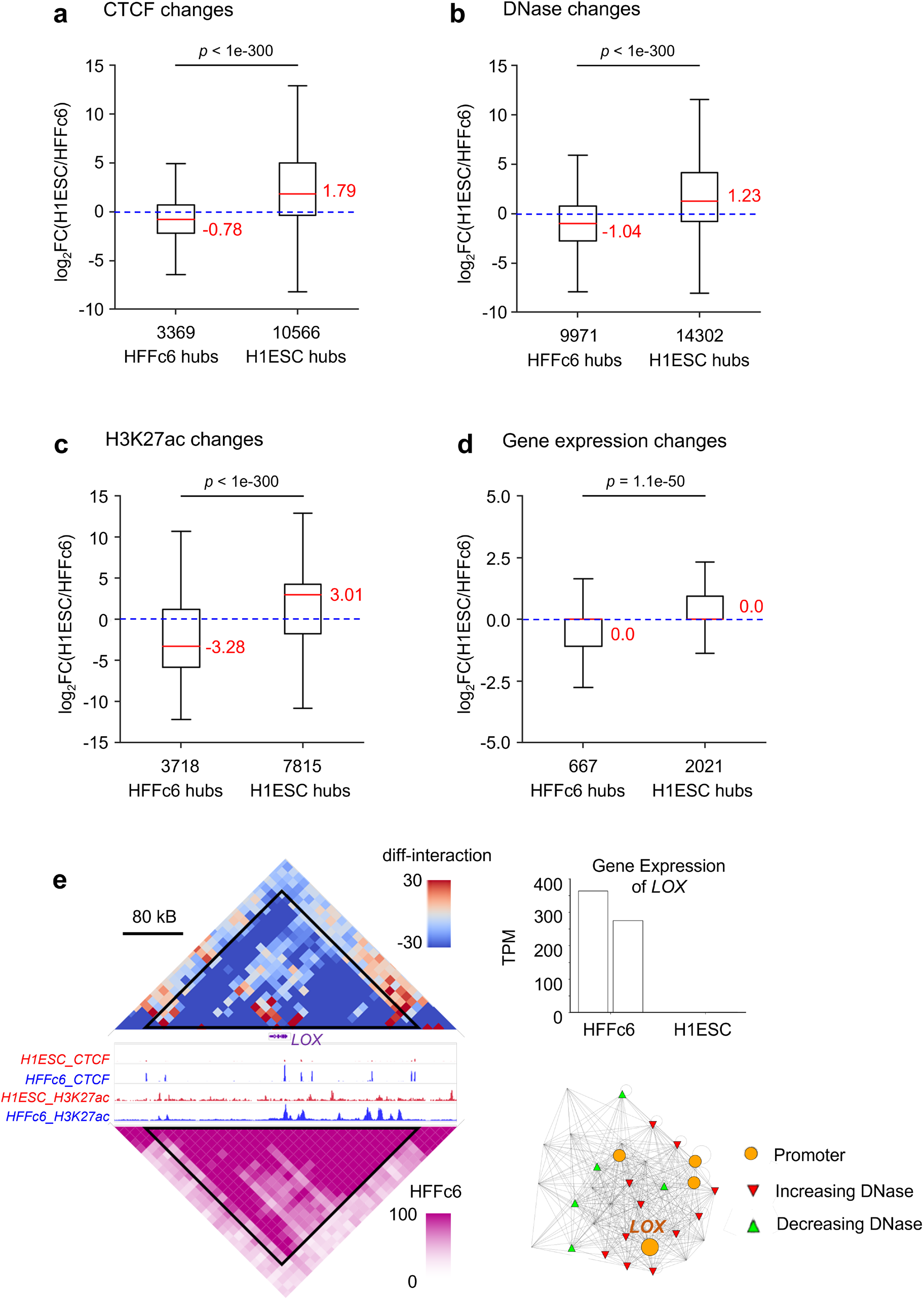
Cell-type-specific hubs exhibit concordant changes in their functional genomic state. **a**. Boxplots showing the distribution of changes of CTCF binding on hub-anchors. CTCF peaks from both cell-types were combined, and the changes of CTCF binding on the combined set of peaks were evaluated. **b**. Boxplots showing the distribution of changes of chromatin accessibility on hub-anchors. DNase hypersensitive sites (DHSs) from both cell-types were combined, and the changes of chromatin accessibility on the combined DHSs were evaluated. **c**. Boxplots showing the distribution of changes of histone mark H3K27ac on hub-anchors. H3K27ac peaks from both cell-types were combined, and the changes of H3K27ac level on the combined peaks were evaluated. **d**. Boxplots showing the distribution of expression changes of hub-associated genes. In (**a-d**) and other box plots, the red line in the middle of the box denotes the median, box denotes interquartile range (IQR), and whiskers denote the most extreme data points that are no more than 1.5 × IQR from the edge of the box; the statistical significance was calculated using one-sided Mann–Whitney U test. **e**. A HFFc6-specific hub enclosing the *LOX* locus and its network visualization. The diamond graph on the top and bottom left showed the differences in chromatin interactions (H1-ESC - HFFc6), and the chromatin interactions in the HFFc6 cells. The highlighted triangle represented the hub. The gene structures, CTCF and H3K27ac tracks were displayed in the middle. Genomic and color scales were shown on the top. The expression pattern of *LOX*, two replicates for each cell-type, was displayed on the top half of the right side. The hub-associated network plot was shown on the bottom half of the right side. Nodes represented genomic elements and edges interactions that changed in the desired direction. Only the subset of nodes and edges in the associated network community belonging to the hub was included. The networks can embed annotation. The circles represented promoters and the triangles represented H1-ESC-specific (green) and HFFc6-specific (red) DNase hypersensitive sites.

Associating a gene with a hub when its promoter overlapped with a hub anchor and using the transcriptome in H1ESC cells from ENCODE^27, 28^ and HFFc6 cells from the 4DN network^19, 29, 30^ (Table S1), we found genes associated with H1-ESC-specific hubs and with HFFc6-specific hubs exhibited significantly different directions of expression changes (Fig. 2d). An example of HFFc6-specific hub formed by intra-interactions within a single hub anchor (i.e., the triangle shape) and the associated differentially expressed gene were shown in Fig. 2e. Examples of H1ESC-specific hub formed with a single hub anchor and cell-type-specific hubs formed by hub-anchor pairs were shown in Fig. S3. Taken together, these analyses demonstrated the biological relevance of identified differential hubs, whose anchors exhibited concordant changes in chromatin states and gene expressions.

### Hubs harbor clustering of transcriptional regulator bindings specific to the same cell type

We further asked if the binding sites of transcriptional regulators that displayed changes in chromatin state were uniformly distributed on hub-anchors. For a transcriptional regulator, we identified its cell-type-specific binding sites and evaluated the enrichment level of the clustered sites with nearest-neighbor spacing less than 12.5Kb apart. This choice of spacing was motivated by the stitching parameter used in the identification of SEs^12^ and it was significantly shorter than the average distance between neighboring sites on hubs. Consistent with the result shown in Fig. 2a, we found that cell-type-specific CTCF binding sites were enriched in the hub-anchors of the same cell type and depleted in the opposite cell type. Interestingly, the level of enrichment and depletion were further enhanced when only the clustered sites with the number of sites in a cluster >=2 were tallied (Fig. 3a). We also found H1-ESC-specific master transcription factors SOX2^31, 32^ (Fig. 3b), NANOG^32, 33^ and OCT4^32, 34^ (Fig. S4) exhibited a similar trend. These observations suggested a connection of hubs with SEs. Indeed, we found that cell-type-specific SEs were enriched on hub-anchors of the same cell type and depleted in those of the opposite cell type (Fig. 3c). Taken together, we found that cell-type-specific transcriptional regulator bindings tended to exhibit clustering on the hubs of the same cell type, but not on the hubs of the opposite cell type.

**Figure 3.**
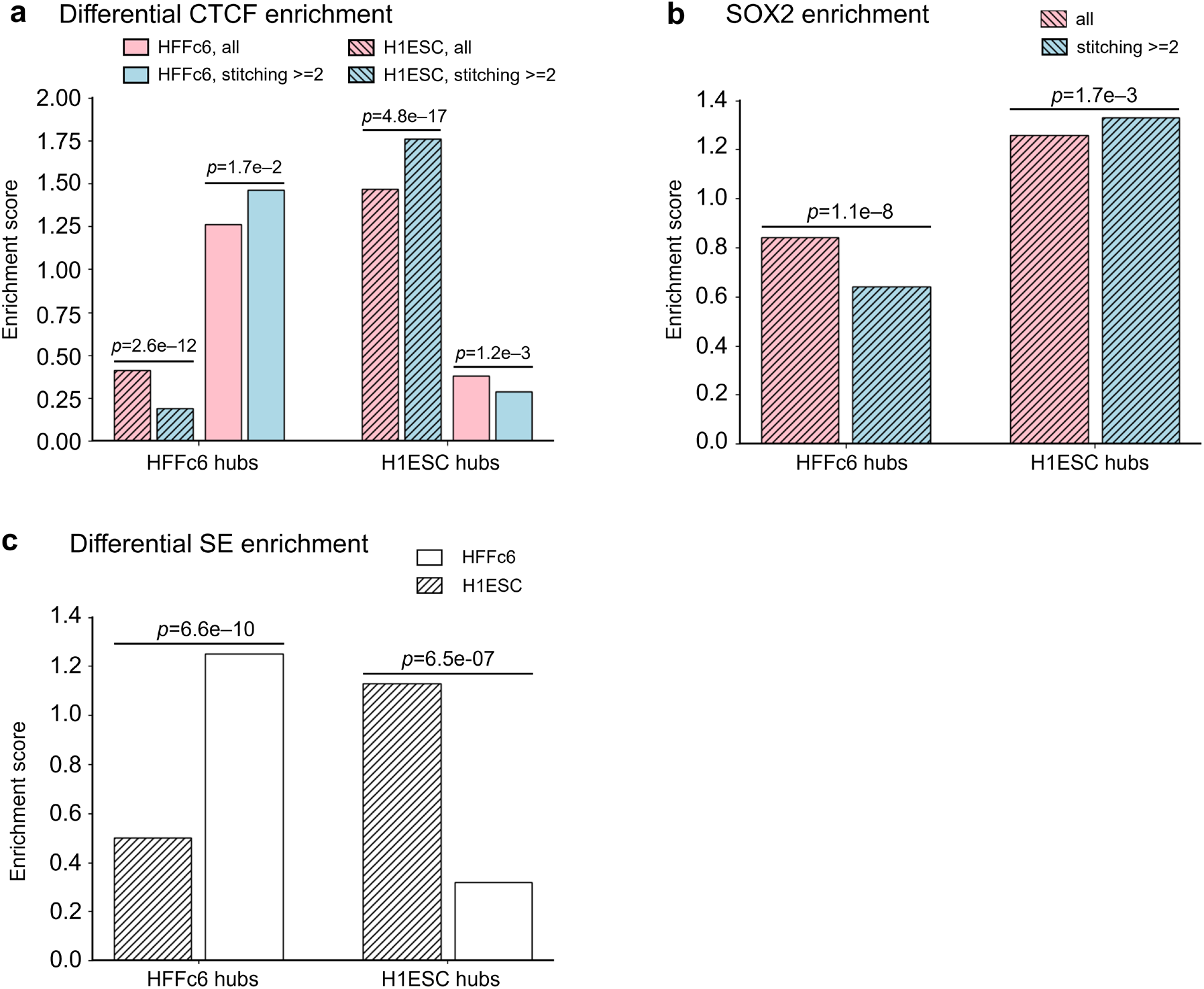
Cell-type-specific hubs harbor clustering of transcriptional regulator binding specific to the same cell type. **a**. Enrichment of cell-type-specific CTCF binding sites on hub-anchors. **b**. Enrichment of SOX2 binding sites on hub-anchors. In (**a-b**) the enrichment score on hub-anchors of HFFc6-specific hubs (left) and H1-ESC-specific hubs (right) was shown for all sites (pink), clustered sites with the number of sites in a cluster >= 2 (blue). Solid and shaded bars represent the enrichment scores calculated using HFFc6-specific and H1-ESC-specific sites, respectively. SOX2 is a H1-ESC-specific transcription factor. The enrichment score was calculated as the ratio of the number of observed occurrences versus the expected, which was evaluated from the genome average. **c**. Enrichment of cell-type-specific super-enhancers on hub-anchors. Statistical significance was determined using one-sided binomial test for (**a**-**c**).

### HiCHub is robust to sequencing depth

To test the effect of sequencing coverage, we used only one technical replicate of Micro-C data from H1-ESC and HFFc6 cells. With less than 250 million reads per library, they are a 10-fold reduction in sequencing depth compared to the full data set (See method for details). Given the lower sequencing coverage, we reduced the significance threshold of hubs to *p*-value < 1E-5 while maintaining the resolution of the contact matrices at 10Kb. We detected 2229 H1-ESC-specific hubs and 1376 HFFc6-specific hubs. In agreement with results obtained from the full Micro-C data set, the hub-anchors of the differential hubs exhibited concordant changes in chromatin states and gene expressions (Fig. S5), enhanced enrichment and depletion of clustering of differential CTCF binding and SEs on hub-anchors (Fig. S6).

## Discussion

In this work we introduced HiCHub, which employed a network-based approach to identify hubs of genomic elements that are in physical proximity in one state and exhibited concordant decreases in their interactions with each other in the other state. In contrast to the existing approaches that focused on the significant changes of individual elements of the contact matrix, HiCHub used the consistency of interaction changes across a set of matrix elements to determine significance. As such, HiCHub is uniquely able to identify sets of concordantly changing chromatin interactions that individually might only exhibit moderate changes, providing an analysis complementary to the prevailing approaches that identify differential loops. Similar ideas have been successfully employed in gene set enrichment analysis^35, 36^, and the analysis of histone modifications^37^ and SEs in the one-dimensional epigenome.

The hub anchors exhibited a concordant pattern of changes in chromatin state, including CTCF binding, open chromatin, and histone mark H3K27ac, indicating their participation in gene regulation. Given our findings on the enrichment of differential SEs on hubs and given the connection of SE with the transcriptional condensate^13^, it is plausible that the identified hubs may reflect the changes in the chromatin architecture associated with the formation and dissolution of condensate. Despite considerable effort in the identification of significant interactions and significant differential interactions, their connection to gene expression remains a challenge. The hub-associated genes exhibit a pattern of expression change concordant with the cell-state specificity of the hubs, facilitating the molecular dissection of specific regulatory circuits, including both the participating regulatory elements and their target gene. Therefore, HiCHub contributes to improving the explanatory power of 3D genome. On the other hand, many genes associated with the hub do not exhibit significant changes in expression (Fig. 2d). Therefore, further refinement in the identification of the target genes of a hub would be desirable and a direction for future studies.

## Method

### HiCHub pipeline

We developed a network approach to systematically compare the differences in chromatin\ interaction between two experimental conditions or cell types/states. To illustrate the procedure, here we use the human H1-ESC and HFFc6 cells to identify H1-ESC-specific hubs as an example. Contact matrices in .hic file format with the same resolution for H1-ESC and HFFc6 cells serve as input. **1)** Data preprocessing. To reduce noise and speed up the calculation, matrix elements whose combined read count from H1-ESC and HFFc6 are less than a cutoff threshold were removed from further consideration. The cutoff threshold is 10 by default and can be set by the user. To enable fair comparison, we then normalize the filtered contact matrices based on the assumption that at any distance the majority of interactions are unchanged. Specifically, for each matrix element, the geometric mean of H1-ESC and HFFc6 interaction values was calculated, the matrix elements were then stratified according to the geometric mean values. For each stratified partition, the sum of the matrix elements belonging to that partition was taken for H1-ESC and HFFc6 cells, respectively. The ratio of the two sums was used as the normalization factor and applied to the HFFc6 contact matrix. This procedure was performed for every partition. After the normalization, the read-count distribution of Log_2_(HFFc6/H1-ESC) for every stratified partition was expected to be centered around zero. **2)** Find the matrix of changed interactions. Normalized matrix elements whose value is higher in H1-ESC cells were selected for the next step. **3)** Network clustering. Chromatin contacts selected in step 2 were used to construct a network using the igraph platform^20^, where the nodes correspond to the anchors (i.e., genomic bin) of contacts and the edges the contacts themselves. Clusters on the network were identified using the community_multilevel algorithm (return-level = 0) with the edge weights set as the original contact matrix element value in H1-ESC cells. This choice ensures that the nodes in the identified community are spatially close in the reference H1-ESC state. **4)** Projection to the genome. The network communities were then projected onto the genome to identify genomic regions as follows. For each network community, each genomic anchor bin was scored by the pagerank value of the corresponding network node. Bins with pagerank values higher than the chromosomal median of pagerank values were designated as eligible and subject to stitching along the chromosome allowing gaps of up to 1 bin, resulting in a collection of stitched regions. For each pair of the regions after the stitching operation (including self), contacts connecting the two regions form either a pyramid or a stripe out of the contact matrix, which serves as a candidate hub. **5)** Significance evaluation. The distribution of the difference of the matrix elements inside each candidate hub was calculated, and a one-sided Wilcoxon signed-rank test was used to evaluate the significance of whether the contacts in the candidate hub collectively changed in the desired direction. The hubs with significance above the preset threshold (default p-value = 1e-5) were considered significant. The identification of HFFc6-specific hubs followed the same procedure. It started with chromatin contacts that exhibited preponderance in HFFc6 cells in step 2) and used the original contact matrix element value in HFFc6 cells for network clustering in step 3).

### Micro-C data set used for HiCHub analysis

The contact matrix of the pooled data in H1-ESC (Accession ID: 4DNFI2TK7L2F) and in HFFc6 cells (Accession ID: 4DNFIPC7P27B) was downloaded from 4DN networks^19, 29, 30^ and used for HiCHub analysis unless specified otherwise (Table S1). The resolution for the two input matrices was chosen to be 10Kb.

### Comparison of CTCF binding on hubs

The CTCF ChIP-Seq peak bed files and normalized count bigwig files were downloaded from the ENCODE consortium^27, 28^ (Table S1). There are 57,383 and 90,470 peaks in H1-ESC and HFFc6 cell, respectively. The peaks called from the two cell-types were merged into a set of consensus peaks using BeEDTools^38^. The read coverage on the consensus peaks from the two cell-types were calculated from the bigwig files using the pyBigWig tool (https://github.com/deeptools/pyBigWig). For each cell type, the read coverage values were then normalized by the total read coverage on all the consensus peak regions. The fold-change of CTCF between the two cell-types on each consensus peak was calculated using the normalized read coverage. The fold-change values were further adjusted by the median of CTCF fold change on all consensus peaks so that the median of log2(fold-change) value after adjustment becomes 0. The distribution of the CTCF fold-change on consensus peaks on cell-type-specific hubs was summarized in boxplots.

### Identification of cell-type-specific CTCF binding sites and their enrichment on hubs

The cell-type-specific CTCF binding sites were identified as those with fold-change > 4 and site coverage from bigwig >= 5 in the cell-type with higher coverage. There were 21,109 and 12,752 sites specific to H1-ESC and HFFc6 cells, respectively. The enrichment score of the cell-type-specific CTCF binding on cell-type-specific hubs is defined as the ratio of the observed number of cell-type-specific CTCF binding versus the expected, which is calculated as the genome-wide density of cell-type-specific CTCF binding times the total length of cell-type-specific hubs. A score greater or smaller than 1 indicates enrichment or depletion, respectively.

### Comparison of chromatin accessibility on hubs

The DNase-Seq peak bed files and normalized count bigwig files were downloaded from the ENCODE consortium^27, 28^ (Table S1). 38,200 and 152,293 DNase hypersensitive sites (DHSs) are called in H1-ESC and HFFc6 cell, respectively Comparison of DHS signal levels on hubs between two cell types was conducted using the same approach as that adopted for CTCF.

### Identification of cell-type-specific chromatin accessibility on hubs and motif analysis

The cell-type-specific DHSs were identified as those with fold-change > 8 and site coverage from bigwig >= 10 in the cell-type with higher coverage. 24,605 and 22,203 DHSs were found specific to H1-ESC and HFFc6 cells, respectively. Cell-type-specific DHSs on the cell-type-specific hubs were identified. The enrichment score of the cell-type-specific DHSs on cell-type-specific hubs was calculated using the same approach as that adopted for CTCF.

### Comparison of histone mark H3K27ac on hubs

H3K27ac peak bed files and normalized-count bigwig files were downloaded from the ENCODE Consortium^27, 28^ (Table S1). Comparison of H3K27ac signal levels on hubs between two cell types was conducted using the same approach as that adopted for CTCF.

### Gene expression analyses

Expression values (TPM) for H1-ESC cells were obtained from the ENCODE consortium (Accession ID: ENCFF321HCT, GEO accession: GSE90225). The expression values (TPM) for HFFc6 cells were obtained from the 4DN consortium (Accession ID: 4DNFI5MR6C3G). Genes whose promoters lie within a hub were considered associated with the hub.

### Enrichment analysis of clustered sites of transcription regulator binding or open chromatin

Clustering of peaks along the genome was carried out by stitching together peaks that are spaced less than 12.5Kb apart. The enrichment score of the clustered peaks on cell-type-specific hubs is defined as the ratio of the observed number of such peaks versus the expected, which was calculated as the genome-wide density of such type of peaks times the total length of cell-type-specific hubs. A score greater or smaller than 1 indicates enrichment or depletion, respectively.

### Enrichment analysis of cell-type-specific super enhancers

Super enhancers (SEs) in H1-ESC and HFFc6 cells were called with the ROSE algorithm^12^ using default parameters applied to the H3K27ac data. The SEs called from the two cell-types were merged into a set of consensus SEs. The normalized read coverage on the consensus SEs from the two cell-types were calculated from the normalized bigwig files. The fold-change of K27ac between the two cell-types on each consensus SE was calculated using the normalized read coverage. Cell-type-specific SEs were identified those with fold-change > 4. 1,083 and 402 Ses were found specific to H1-ESC and HFFc6 cells, respectively. A SE was identified to be associated with a hub if its midpoint overlapped with a hub-anchor.

### Reproducibility of hubs

Cell-type-specific hubs were called with the contact matrix obtained from pooling all data, including both technical and biological replicates, of the Micro-C data set in H1-ESC (Accession ID: 4DNES21D8SP8) and HFFc6 cells (Accession ID:4DNESWST3UBH), respectively. For each pair of individual biological replicates from the two cell types and for each of the cell-type-specific hubs, we calculated the median of log2(fold-change) of matrix elements associated with the hubs. We then used the correlation of these values to evaluate reproducibility. Using the method described above, we evaluated 2 replicate pairs. Replicate-pair 1 were prepared by combining the Micro-C data from technical replicates of bio-replicate 1 of H1-ESC cells (Accession ID: 4DNFI8GM4EL9, 4DNFIMT4PHZ1, 4DNFIBMG8YA3, 4DNFING6ZFDF) and technical replicates of bio-replicate 3 of HFFc6 (Accession ID: 4DNFIV8JNJB8, 4DNFIJM966UR, 4DNFIPKVL9YI, 4DNFIG1ZOVIM, 4DNFIONHB78N, 4DNFIFJL4JIH), respectively. We matched these two because they have similar sequencing coverage. Replicate-pair 2 were prepared by combining the Micro-C data from technical replicates of bio-replicate 2 of H1-ESC cells (Accession ID: 4DNFIULY29IQ, 4DNFI2YHYAJO, 4DNFIXP9MVBU, 4DNFI89L17XY, 4DNFIIYUGYBU) and technical replicates of bio-replicate 2 of HFFc6 (Accession ID: 4DNFIKWV6BY2, 4DNFIZU6ADT1, 4DNFIATCW955, 4DNFIK82YRNM, 4DNFIF5F4HRG), respectively.

### Applying HiCHub to libraries of limited sequencing depth

The following Micro-C libraries were used for input of HiCHub: Technical replicate 3 of bio-replicate 1 (Accession ID: 4DNFIMT4PHZ1) for H1-ESC cells and technical replicate 4 of bio-replicate 2 (Accession ID: 4DNFIK82YRNM) for HFFc6 cells. They were chosen because of matching sequencing depth: 284 million for H1-ESC and 298 million for HFFc6. In contrast, the sequencing depth for the full pooled data was 3.22 billion for H1-ESC and 5.86 billion for HFFc6.

## Data availability

The high-throughput sequencing data used in the study are summarized in the Supplementary Table 1.

## Code availability

The source code for HiCHub is freely available at https://github.com/WeiqunPengLab/HiCHub.

## Acknowledgements

This study is supported in-part by grants from the NIH (AI121080 and AI139874 to H.-H.X. and W.P., AI112579 to H.-H.X.), the GWU CDRF to W. P. and the Veteran Affairs BLR&D Merit Review Program (BX002903) to H.-H.X.

## Conflict of Interest

none declared.

## Notes

### Competing Interest Statement

The authors have declared no competing interest.

## References

1. Yu, M. & Ren, B. The Three-Dimensional Organization of Mammalian Genomes. Annu Rev Cell Dev Biol 33, 265–289 (2017).

2. Zheng, H. & Xie, W. The role of 3D genome organization in development and cell differentiation. Nat Rev Mol Cell Biol 20, 535–550 (2019).

3. Hu, G. et al. Transformation of Accessible Chromatin and 3D Nucleome Underlies Lineage Commitment of Early T Cells. Immunity 48, 227–242 e228 (2018).

4. Shan, Q. et al. Tcf1 preprograms the mobilization of glycolysis in central memory CD8(+) T cells during recall responses. Nat Immunol 23, 386–398 (2022).

5. Spielmann, M., Lupianez, D.G. & Mundlos, S. Structural variation in the 3D genome. Nature reviews. Genetics 19, 453–467 (2018).

6. Dixon, J.R. et al. Integrative detection and analysis of structural variation in cancer genomes. Nat Genet 50, 1388–1398 (2018).

7. Lun, A.T.L. & Smyth, G.K. diffHic: a Bioconductor package to detect differential genomic interactions in Hi-C data. BMC Bioinformatics 16, 258 (2015).

8. Djekidel, M.N., Chen, Y. & Zhang, M.Q. FIND: difFerential chromatin INteractions Detection using a spatial Poisson process. Genome Research 28, 412–422 (2018).

9. Stansfield, J.C., Cresswell, K.G. & Dozmorov, M.G. multiHiCcompare: joint normalization and comparative analysis of complex Hi-C experiments. Bioinformatics 35, 2916–2923 (2019).

10. Ardakany, A.R., Ay, F. & Lonardi, S. Selfish: discovery of differential chromatin interactions via a self-similarity measure. Bioinformatics 35, i145–i153 (2019).

11. Cook, K.B., Hristov, B.H., Le Roch, K.G., Vert, J.P. & Noble, W.S. Measuring significant changes in chromatin conformation with ACCOST. Nucleic Acids Res 48, 2303–2311 (2020).

12. Whyte, W.A. et al. Master transcription factors and mediator establish super-enhancers at key cell identity genes. Cell 153, 307–319 (2013).

13. Sabari, B.R. et al. Coactivator condensation at super-enhancers links phase separation and gene control. Science 361 (2018).

14. Ing-Simmons, E. et al. Spatial enhancer clustering and regulation of enhancer-proximal genes by cohesin. Genome Res 25, 504–513 (2015).

15. Madsen, J.G.S. et al. Highly interconnected enhancer communities control lineage-determining genes in human mesenchymal stem cells. Nat Genet 52, 1227–1238 (2020).

16. Beagrie, R.A. et al. Complex multi-enhancer contacts captured by genome architecture mapping. Nature 543, 519–524 (2017).

17. Petrovic, J. et al. Oncogenic Notch Promotes Long-Range Regulatory Interactions within Hyperconnected 3D Cliques. Molecular Cell 73, 1174-1190.e1112 (2019).

18. Akgol Oksuz, B. et al. Systematic evaluation of chromosome conformation capture assays. Nat Methods 18, 1046–1055 (2021).

19. Krietenstein, N. et al. Ultrastructural Details of Mammalian Chromosome Architecture. Mol Cell 78, 554–565 e557 (2020).

20. Csárdi, G. & Nepusz, T. The igraph software package for complex network research. 2006; 2006.

21. Braccioli, L. & de Wit, E. CTCF: a Swiss-army knife for genome organization and transcription regulation. Essays Biochem 63, 157–165 (2019).

22. Ghirlando, R. & Felsenfeld, G. CTCF: making the right connections. Genes Dev 30, 881–891 (2016).

23. Arzate-Mejía, R.G., Recillas-Targa, F. & Corces, V.G. Developing in 3D: the role of CTCF in cell differentiation. Development 145 (2018).

24. Ren, G. et al. CTCF-Mediated Enhancer-Promoter Interaction Is a Critical Regulator of Cell-to-Cell Variation of Gene Expression. Mol Cell 67, 1049-1058.e1046 (2017).

25. Huang, J. et al. Dissecting super-enhancer hierarchy based on chromatin interactions. Nature Communications 9, 943 (2018).

26. Kai, Y. et al. Predicting CTCF-mediated chromatin interactions by integrating genomic and epigenomic features. Nature Communications 9, 4221 (2018).

27. Consortium, E.P. An integrated encyclopedia of DNA elements in the human genome. Nature 489, 57–74 (2012).

28. Davis, C.A. et al. The Encyclopedia of DNA elements (ENCODE): data portal update. Nucleic Acids Res 46, D794–D801 (2018).

29. Dekker, J. et al. The 4D nucleome project. Nature 549, 219–226 (2017).

30. Reiff, S.B. et al. The 4D Nucleome Data Portal: a resource for searching and visualizing curated nucleomics data. bioRxiv, 2021.2010.2014.464435 (2021).

31. Zhou, C. et al. Comprehensive profiling reveals mechanisms of SOX2-mediated cell fate specification in human ESCs and NPCs. Cell Res 26, 171–189 (2016).

32. Zheng, R. et al. Cistrome Data Browser: expanded datasets and new tools for gene regulatory analysis. Nucleic Acids Res 47, D729–D735 (2019).

33. Lister, R. et al. Human DNA methylomes at base resolution show widespread epigenomic differences. Nature 462, 315–322 (2009).

34. Ji, X. et al. 3D Chromosome Regulatory Landscape of Human Pluripotent Cells. Cell stem cell 18, 262–275 (2016).

35. Subramanian, A. et al. Gene set enrichment analysis: A knowledge-based approach for interpreting genome-wide expression profiles. Proceedings of the National Academy of Sciences 102, 15545–15550 (2005).

36. Mootha, V.K. et al. PGC-1α-responsive genes involved in oxidative phosphorylation are coordinately downregulated in human diabetes. Nature Genetics 34, 267–273 (2003).

37. Zang, C. et al. A clustering approach for identification of enriched domains from histone modification ChIP-Seq data. Bioinformatics (Oxford, England) 25, 1952–1958 (2009).

38. Quinlan, A.R. & Hall, I.M. BEDTools: a flexible suite of utilities for comparing genomic features. Bioinformatics 26, 841–842 (2010).

