## Supplemental Material for "HiCHub: A Network-Based Approach to Identify Domains of Differential Interactions from 3D Genome Data"

Supplementary data include 6 supplementary figures and 1 supplementary table.

#### Supplementary Figure 1

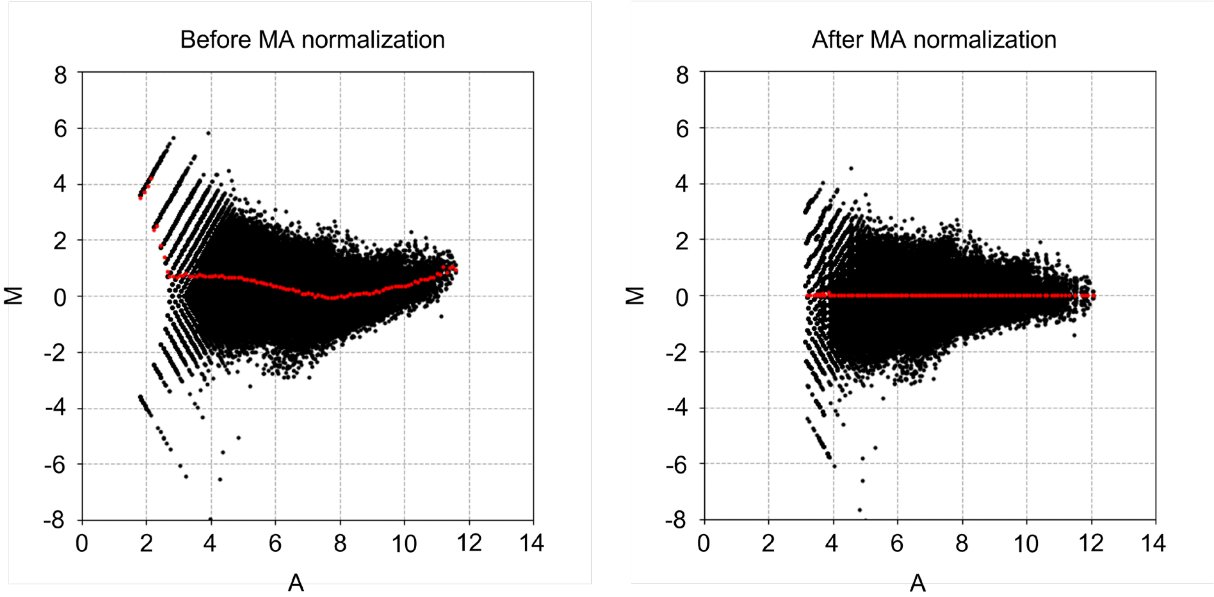

**Supplementary Figure 1. Illustration of the MA normalization.** Shown are the MA plots of chromatin interactions on chromosome 1 before (left) and after (right) MA normalization. The x- and y-axis denote the average (A), (log2 (H1-ESC) + log2 (HFFc6))/2, and mass (M), log2 (H1-ESC/HFFC6), respectively. The red dotted curve represents the average of M values at a given A value.

### Supplementary Figure 2

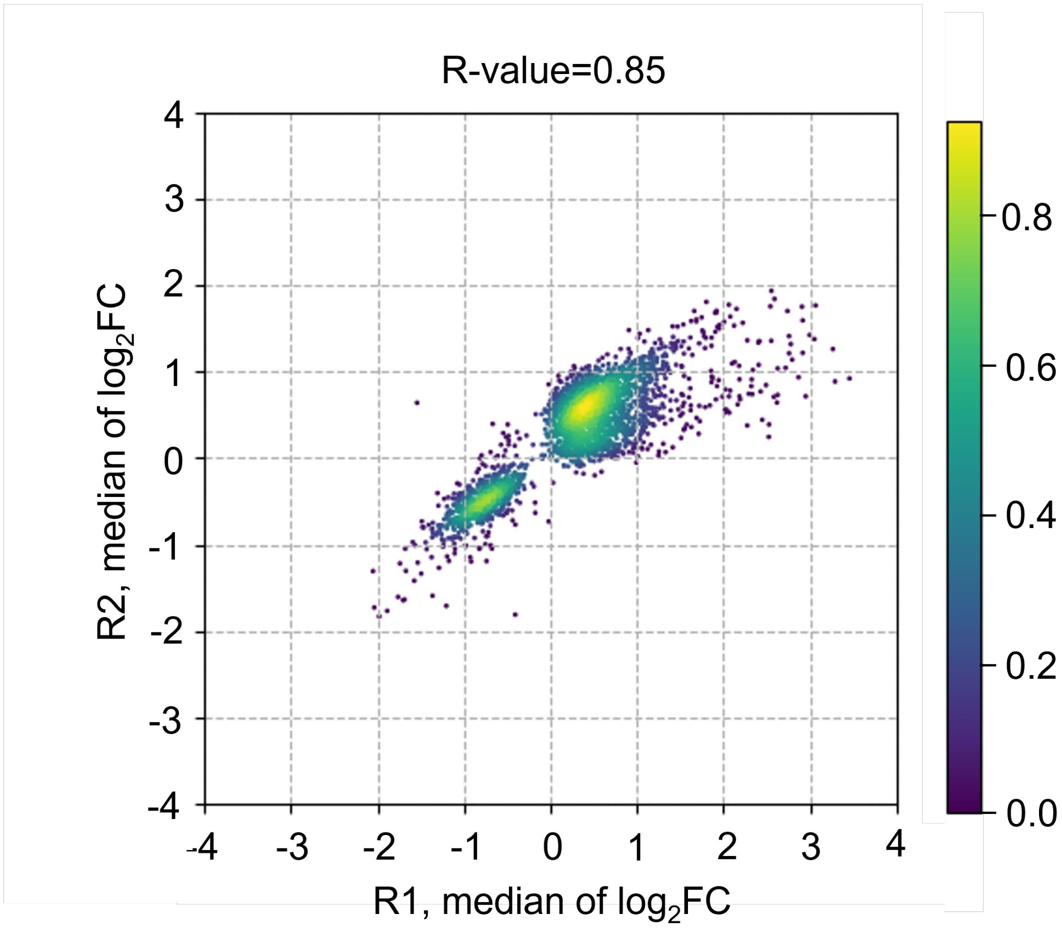

**Supplementary Figure 2. Reproducibility analysis of hubs using biological replicates.** Two pairs of biological replicates of Micro-C data were used to evaluate interaction changes on hubs called from the pooled Micro-C data (see Methods for details). For each pair of biological replicates and for each hub, the median value of log2(fold-change) of matrix elements were calculated and were used for the scatter plot.

#### Supplementary Figure 3

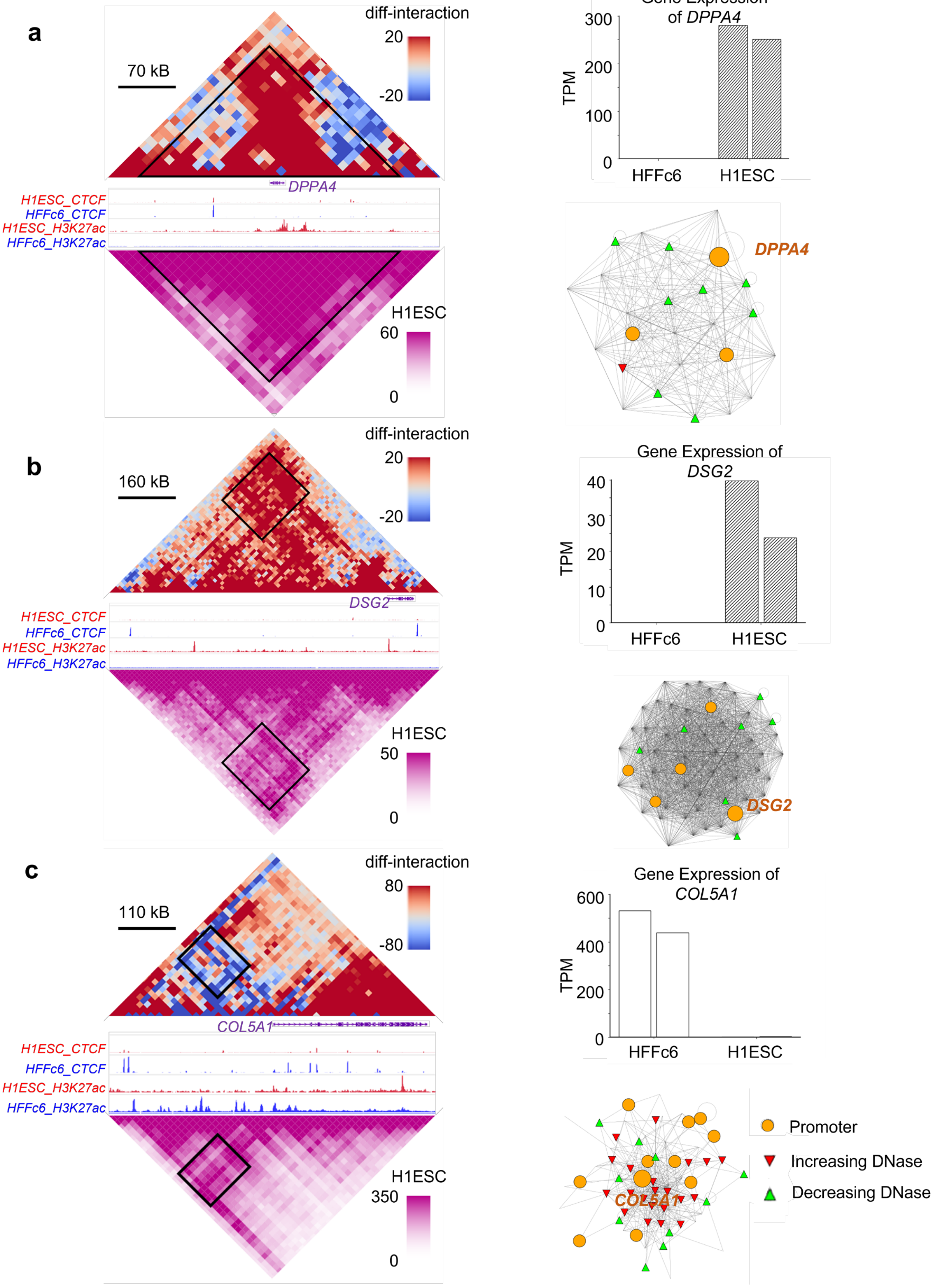

**Supplementary Figure 3. Examples of hubs with two hub-anchors.**

**a.** A H1-ESC-specific hub enclosing the *DPPA4* locus.

**b.** A H1-ESC-specific hub with two hub-anchors associated with the *DSG2* locus.

**c.** A HFFc6-specific hub with two hub-anchors associated with the *COL5A1* loci.

The diamond graph on the top and bottom left showed the differences in chromatin interactions (H1-ESC - HFFc6), and the chromatin interactions in the H1-ESC (**a, b**) or HFFc6 (**c**) cells, respectively. The highlighted triangle or rectangle represented the hub. The gene structures, CTCF and H3K27ac tracks were displayed in the middle. Genomic and color scales were shown on the top. The expression pattern of the hub-associated differentially expressed gene (two replicates for each cell-type) and the hub-associated network plot were shown on the right part of each panel. Nodes represented genomic elements and edges interactions that changed in the desired direction. Only the subset of nodes and edges in the associated network community belonging to the hub was included. The networks can embed annotation. The circles represented promoters and the triangles represented H1-ESC-specific (green) and HFFc6-specific (red) DNase hypersensitive sites.

#### Supplementary Figure 4

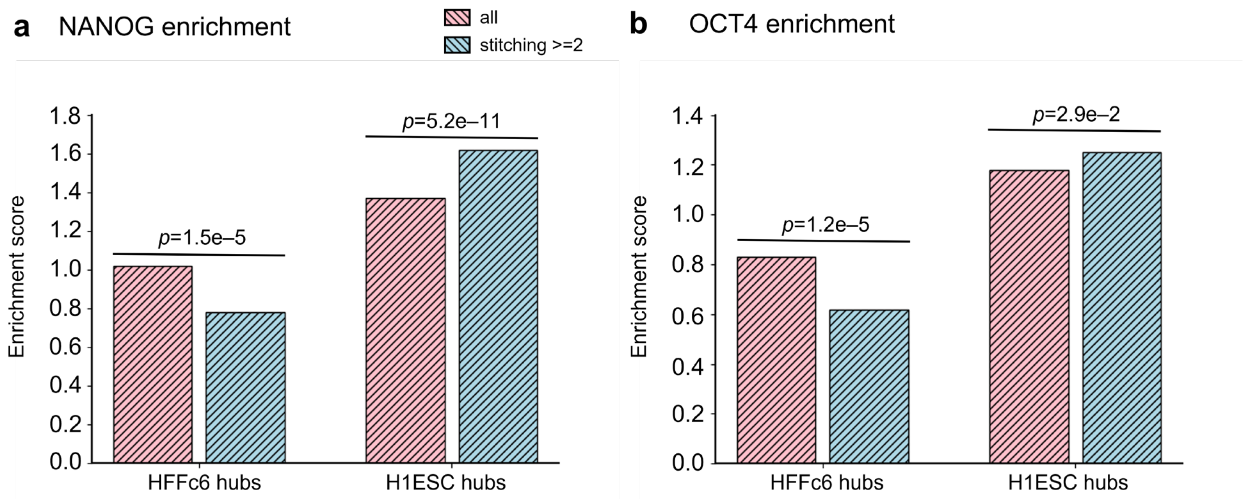

**Supplementary Figure 4. Enrichment of NANOG (a) and OCT4 (b) binding sites on hubs.** The enrichment score on hub-anchors of HFFc6-specific hubs (left) and H1-ESC-specific hubs (right) was shown for all sites (pink), clustered sites with the number of sites in a cluster >= 2 (blue). The enrichment score was calculated as the ratio of the number of observed occurrences versus the expected, which was evaluated from the genome average.  Statistical significance was determined using one-sided binomial test.

#### Supplementary Figure 5

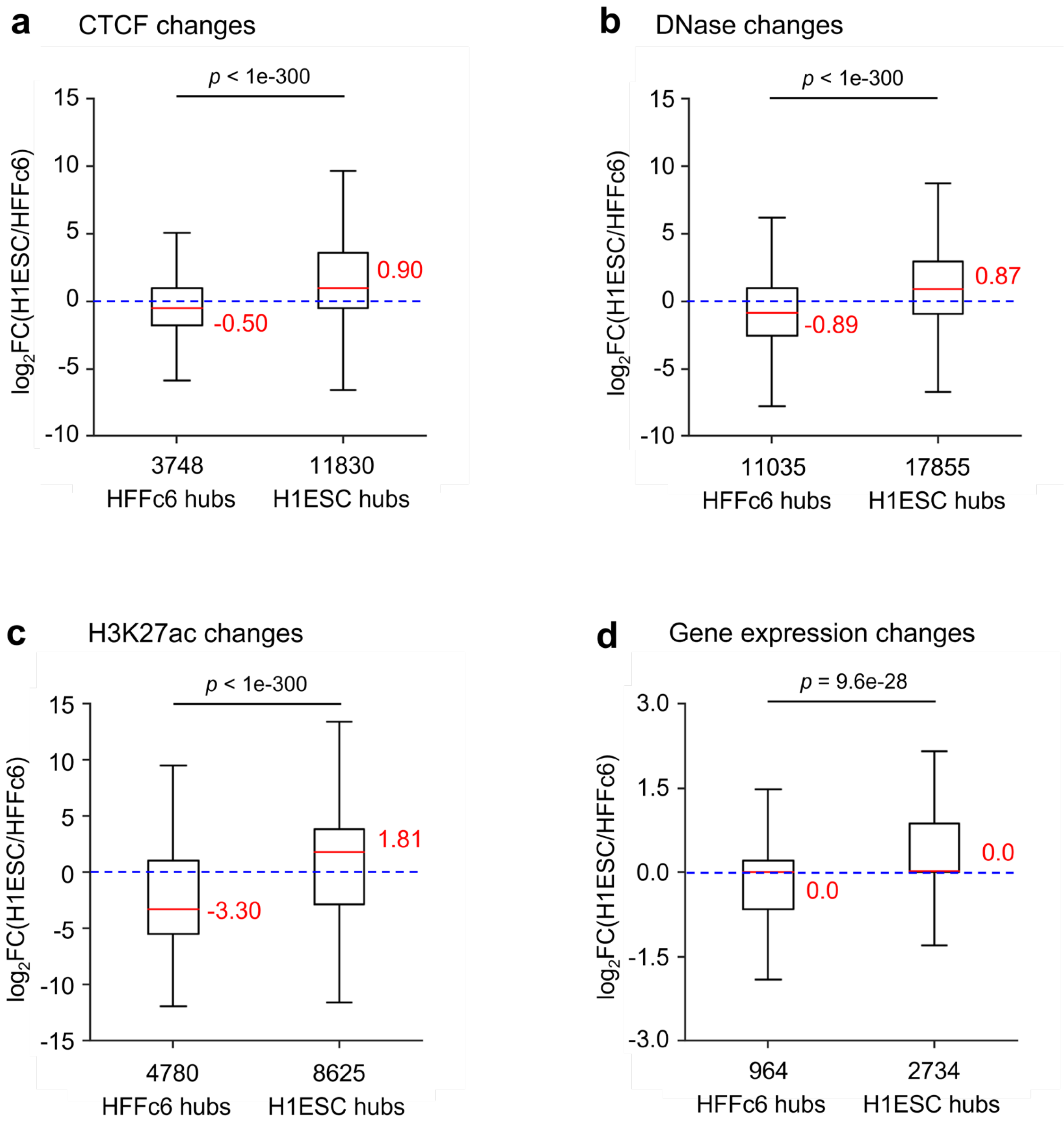

**Supplementary Figure 5. Hubs identified using libraries with low sequencing coverage exhibit concordant changes in their functional genomic state.**

**a.** Boxplots showing the distribution of changes of CTCF binding on hub-anchors. CTCF peaks from both cell-types were combined, and the changes of CTCF binding on the combined set of peaks were evaluated.

**b.** Boxplots showing the distribution of changes of chromatin accessibility on hub-anchors. DNase hypersensitive sites (DHSs) from both cell-types were combined, and the changes of chromatin accessibility on the combined DHSs were evaluated.

**c.** Boxplots showing the distribution of changes of histone mark H3K27ac on hub-anchors. H3K27ac peaks from both cell-types were combined, and the changes of H3K27ac level on the combined peaks were evaluated.

**d.** Boxplots showing the distribution of expression changes of hub-associated genes.

Supplementary Figure 6

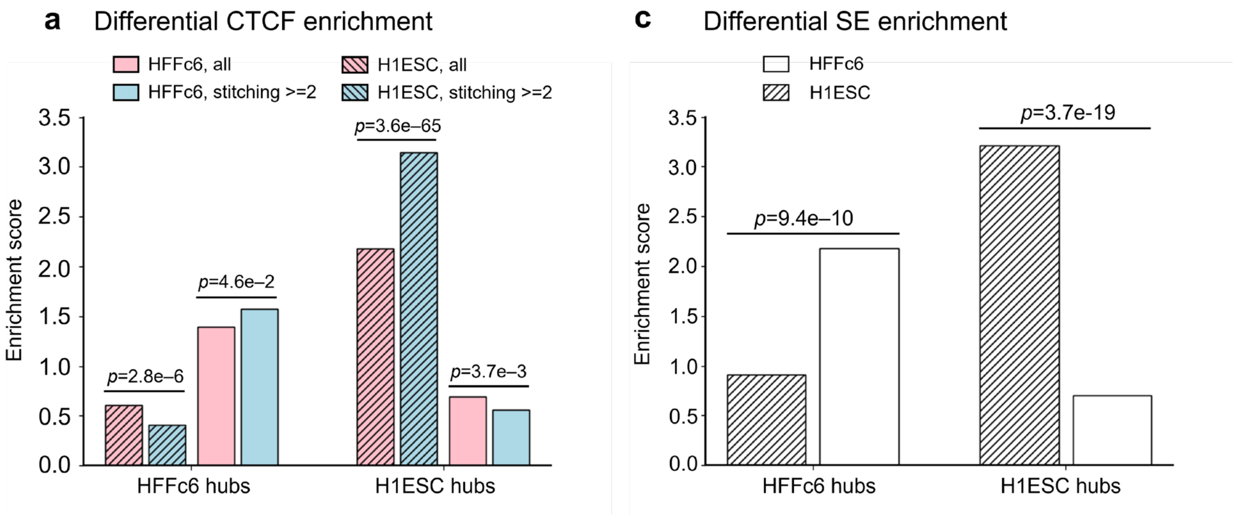

**Supplementary Figure 6**. **Hubs identified using libraries with low sequencing coverage harbor clustering of transcriptional regulator binding specific to the same cell type.**

**a.** Enrichment of cell-type-specific CTCF binding sites on hub-anchors. The enrichment score on hub-anchors of HFFc6-specific hubs (left) and H1-ESC-specific hubs (right) was shown for all sites (pink), clustered sites with the number of sites in a cluster >= 2 (blue). Solid and shaded bars represent the enrichment scores calculated using HFFc6-specific and H1-ESC-specific sites, respectively. The enrichment score was calculated as the ratio of the number of observed occurrences versus the expected, which was evaluated from the genome average.

**b.** Enrichment of cell-type-specific super-enhancers on hub-anchors.

Statistical significance was determined using one-sided binomial test for both (**a**) and (**b**).

### Supplementary Table S1

| Cell type | Experiments | Source | Experiment number | Processed files | Processed number | PMID | GEO |
|---|---|---|---|---|---|---|---|
| H1HESC | Micro-C | 4DN | 4DNES21D8SP8 | Contact matrix (hic) | 4DNFI2TK7L2F | 32213324 | |
| | polyA+ RNA-seq | ENCODE | ENCSR000COU | gene quatifications (tsv) | ENCFF321HCT | 32584815 | GSE90225 |
| | CTCF | ENCODE | ENCSR000BNH | narrow peaks (bed) | ENCFF023LAA | 32657410 | GSM803419 |
| | H3K27ac | Cistrome db | 867 | narrow peaks (bed) | 867 | 19829295 | GSM466732 |
| | | | | normalized counts (bigwig) | | | |
| | DNase | ENCODE | ENCSR000EMU | narrow peaks (bed) | ENCFF983UCL | 27127239 | GSM736582 |
| | | | | normalized counts (bigwig) | ENCFF573NKX | | |
| | SOX2 | Cistrome db | 67318 | narrow peaks (bed) | 67318 | 26809499 | GSM1701825 |
| | OCT4 | Cistrome db | 58494 | narrow peaks (bed) | 58494 | 26686465 | GSM1705267 |
| | NANOG | Cistrome db | 4933 | narrow peaks (bed) | 4933 | 19829295 | GSM456572 |
| HFFc6 | Micro-C | 4DN | 4DNESWST3UBH | Contact matrix (hic) | 4DNFIPC7P27B | 32213324 | |
| | RNA-seq | 4DN | 4DNESFH3EHTU | gene expression (tsv) | 4DNFI5MR6C3G | 28905911 | |
| | CTCF | ENCODE | ENCSR163ULN | narrow peaks (bed) | ENCFF005CJI | 22955616 | GSE175027 |
| | H3K27ac | Cistrome DB | 62055 | narrow peaks (bed) | 62055 | 27127239 | GSM2066623 |
| | | | | normalized counts (bigwig) | | | |
| | DNase | ENCODE | ENCSR672EWY | narrow peaks (bed) | ENCFF462ILY | 22955616 | GSE170964 |
| | | | | normalized counts (bigwig) | ENCFF623ZIV | | |
